# Genetic map of regional sulcal morphology in the human brain

**DOI:** 10.1101/2021.10.13.463489

**Authors:** Benjamin B. Sun, Stephanie J. Loomis, Fabrizio Pizzagalli, Natalia Shatokhina, Jodie N. Painter, Christopher N. Foley, Biogen Biobank Team, Megan E. Jensen, Donald G. McLaren, Sai Spandana Chintapalli, Alyssa H. Zhu, Daniel Dixon, Tasfiya Islam, Iyad Ba Gari, Heiko Runz, Sarah E. Medland, Paul M. Thompson, Neda Jahanshad, Christopher D. Whelan

## Abstract

The human brain is a complex organ underlying many cognitive and physiological processes, affected by a wide range of diseases. Genetic associations with macroscopic brain structure are emerging, providing insights into genetic sources of brain variability and risk for functional impairments and disease. However, specific associations with measures of local brain folding, associated with both brain development and decline, remain under-explored. Here we carried out detailed large-scale genome-wide associations of regional brain cortical sulcal measures derived from magnetic resonance imaging data of 40,169 individuals in the UK Biobank. Combining both genotyping and whole-exome sequencing data (∼12 million variants), we discovered 388 regional brain folding associations across 77 genetic loci at *p*<5×10^−8^, which replicated at *p*<0.05. We found genes in associated loci to be independently enriched for expression in the cerebral cortex, neuronal development processes and differential regulation in early brain development. We integrated coding associations and brain eQTLs to refine genes for various loci and demonstrated shared signal in the pleiotropic *KCNK2* locus with a cortex-specific *KCNK2* eQTL. Genetic correlations with neuropsychiatric conditions highlighted emerging patterns across distinct sulcal parameters and related phenotypes. We provide an interactive 3D visualisation of our summary associations, making complex association patterns easier to interpret, and emphasising the added resolution of regional brain analyses compared to global brain measures. Our results offer new insights into the genetic architecture underpinning brain folding and provide a resource to the wider scientific community for studies of pathways driving brain folding and their role in health and disease.

## Main

Human brain structure and function are complex drivers of basic and higher cognitive processes, which vary between individuals and in numerous neurological, psychiatric and cognitive disorders. Structural magnetic resonance imaging (MRI) scans provide a reliable, non-invasive measure of brain structure and are widely used in research and clinical settings. Genetic variants influencing brain structure and function are important to identify, as they can help uncover pathophysiological pathways involved in heritable brain diseases and prioritize novel targets for drug development. Several large-scale genome-wide association studies (GWAS) have identified hundreds of genetic influences on variations in brain structure and function^1-3^ revealing novel insights into processes guiding brain development, and highlighting potential shared genetic aetiologies with neurodegenerative and psychiatric conditions^4,5^.

To date, most neuroimaging GWAS have focused on broad, macroscale anatomical features such as subcortical volume, cortical thickness and white matter microstructure^6^. Anomalies of cortical gyrification - the folding of the cerebral cortex into its characteristic concave sulci (fissures) and convex gyri (ridges) - contribute to many neurodevelopmental and neuropsychiatric conditions^7,8^, but the genetic underpinnings of gyrification remain relatively understudied^9^. Sulcal characteristics and folding patterns are altered across a range of neurodevelopmental disorders, from cortical dysplasias^10^ to neurogenetic syndromes^11^, and radiologists often use sulcal widening as an early indicator of atrophy in degenerative diseases^12^, as it offers a clear and sensitive biomarker of disease progression^13,14^. Recent neuroimaging genetics investigations have broadened in scale and scope, examining specific sulcal measures across the full brain^15,16^, but without evaluating the reliability of the measures at scale across MRI scanning protocols. Using four independent datasets, we recently outlined a range of heritable sulcal measures that can be reliably quantified at high resolution across the whole brain, irrespective of MRI platform or acquisition parameters^17^.

Here we conducted a comprehensive genome-wide analysis of regional sulcal shape parameters, extracted from the multi-centre brain MRI scans of 40,169 participants in the UK Biobank. To discover rare and common genetic variants influencing cortical gyrification, we conducted GWAS and exome-wide analysis of a total of 450 sulcal parameters^17^. Sulcal shape descriptors, comprising length, mean depth, width, and surface area, were extracted from a discovery cohort of 26,530 individuals of European ancestry and a replication cohort of 13,639 individuals. After mapping the genetic architecture of regional sulcal measures across the cortex, we highlight putative biological and developmental pathway involvement as well as links to neuropsychiatric conditions. Finally, we provide a portal to interactively visualise our results in 3D (https://enigma-brain.org/sulci-browser), demonstrating varying complex patterns of associations, to help inform future investigations of human cortical morphology.

## Results

Regional brain sulcal measurements (4 shape parameters: length, width, mean depth and surface area), regional delineations, and phenotype nomenclature are summarised in **Supplementary Table 1 and Figure 1a**. We determined the overall clustering of the high-dimensional phenotypes with *t*-SNE and found that width parameter phenotypes formed a distinct cluster compared to the other three shape parameters (**Extended Data Figure 1**). Notably, the t-SNE representation retains broad brain lobe topology for the width parameter phenotypes in particular (**Figure 1b)**.

**Figure 1.**
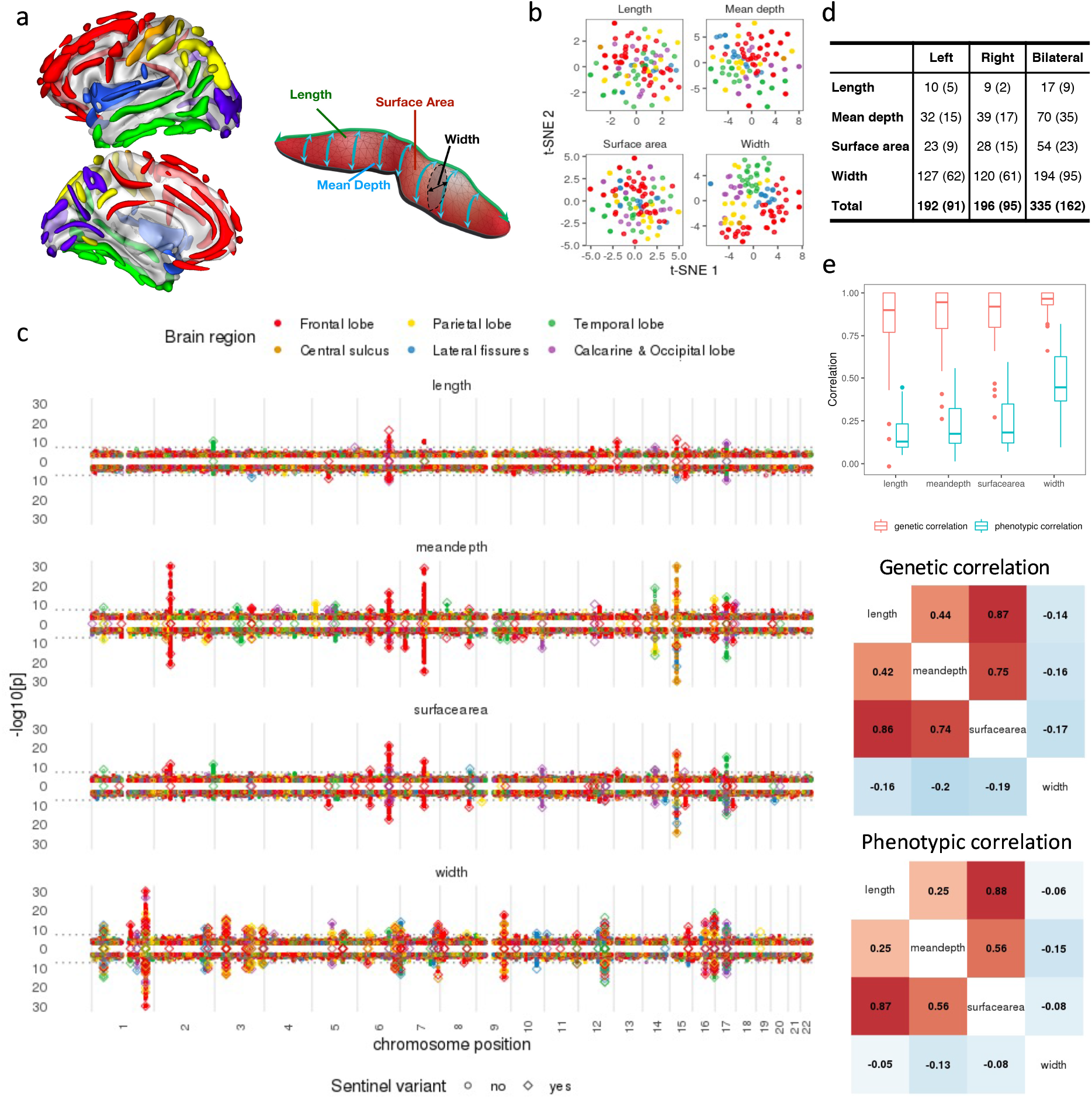
Summary of brain sulcal association results. **(a) Schematic of brain sulcal folds and shape parameters**. Brain region legend corresponds to colours in figures a-c. **(b) t-SNE of regional brain sulcal measures for each shape parameter. (c) GWAS association results by shape parameters and side**. Diamonds indicate lead (sentinel) associations that replicated (*p*<0.05). Points above 0 in the y-axis in each plot refers to associations with left sided sulcal measures, below 0 with right sided measures. Diamonds along 0 is the y-axis indicate lead associations for bilateral sulcal measures. Dashed horizonal line indicate GWAS significance threshold (*p*=5×10^−8^). (**d) Summary of number of associations by side and shape parameters. (e) Top: genetic and phenotypic correlation between left and right sides. Middle: Genetic correlation between shape parameters. Bottom: Phenotypic correlation between shape parameters**. Middle and bottom: left hemisphere correlations in upper triangle, right hemisphere correlations in lower triangle.

### Genetic architecture of regional brain sulcal folds

We conducted GWAS of 450 regional brain sulcal measurements separately for 11.9 million combined imputed and whole-exome sequenced variants in UKB participants divided into a discovery cohort (n=26,530) and a replication cohort (n=13,639) (**Methods, Extended Data Figure 2)**.

At a significance threshold of *p*<2×10^−10^ which accounts for the effective number of independent sulcal measures analysed (**Methods**) **-** we found, and replicated at *p*<0.05, a total of 186 specific sulci parameter associations (for at least one hemisphere) across 41 genetic loci (388 associations across 77 loci at *p*<5×10^−8^) (**Figure 1c and 1d, Extended Data Figure 3, Supplementary Table 2**). We also performed GWAS on bilateral sulcal measurements (averaging values from left and right brain hemispheres) and found a total of 162 replicated associations across 47 loci at *p*<2×10^−10^ (335 associations across 107 loci at *p*<5×10^−8^), where 6 (across 3 loci) and 108 additional associations (across 42 loci) were also found at *p*<2×10^−10^ and *p*<5×10^−8^ respectively (**Supplementary Table 3**). Genomic inflation was well controlled (median λgc=1.02, range: 0.99-1.07). We found an inverse relationship between effect sizes and minor allele frequency (MAF) (**Extended Data Figure 4**), in line with other disease and intermediate trait results, and consistent with variants with strong effects are deleterious and rarer.

We found a similar number of associations for left and right hemispheres. Approximately two-thirds of associations were with sulcal width, followed by mean depth, surface area and length measures, in line with their heritability estimates^17^ **(Figure 1c and 1d, Extended Data Figure 5)**. Length measures accounted for the lowest proportion of associations (<5%, **Figure 1b)**, consistent with length having the lowest heritability (**Extended Data Figure 5)**, especially after adjusting for intra-cranial volume^17^. Comparing the absolute (all effects across hemispheres were consistent in direction) Z-scores of lead associations across hemispheres (left, right and bilateral, **Extended Data Figure 6**), we found no significant difference between left and right hemisphere (paired t-test *p*=0.25), whilst bilateral associations tended to exhibit stronger associations (mean abs(Z-score) 1.00 higher *vs* right, *p*=4.1×10^−96^ and 0.92 higher *vs* left, *p*=1.8×10^−82^), consistent with their heritability estimates (**Extended Data Figure 5 and 6**).

Some genetic loci exhibited highly pleiotropic associations across multiple brain regions; for example, 10 genetic loci were associated with 10 or more sulcal measures, showing different association patterns across shape parameters. Notably, the chr1:215Mb (near *KCNK2*) and chr12:106Mb (12q23.3, *NUAK1*) regions were associated with 23 and 22 width measures respectively across multiple brain regions; the chr16:87Mb region (6q24.2, near *C16orf95*) was associated with 16 width measures across multiple brain regions, 4 mean depth and 1 surface area measures mostly in the frontal lobe; the chr17:47Mb region (17q21.31, containing *MAPT* and *KANSL1*) was associated with 16 width, 9 surface area, 6 mean depth and 2 length measures mostly in the temporal and calcarine-occipital regions; whilst chr6:126Mb region (6q22.32, containing *CENPW*) was associated with 9 surface area, 4 length, 4 mean depth and 2 width measures - mostly in the frontal and calcarine-occipital regions (**Figure 1c, Supplementary Table 2)**.

We cross-referenced the lead variants and their proxies (r^2^>0.8) for significant (*p*<5×10^−8^) associations in previous related brain imaging studies (**Supplementary Information**) in the GWAS Catalog^6^ (LD proxy r^2^>0.8, +/- 500Kb around the lead variant). We found 56 of the 119 loci (77 for left/right hemisphere and 42 bilateral measures, *p*<5×10^−8^) to be associated with any brain imaging phenotype (mostly consisting of brain volume, surface area and white matter microstructure) including the 10 highly pleiotropic genetic loci, many of which (e.g. *CENPW* containing locus (6q22.32), *MAPT-KANSL1* containing locus (17q21.31), *C16orf95* locus (6q24.2), *NUAK1* locus (12q23.3), chr2:65Mb (2p14), chr15:40Mb (15q14), chr14:59Mb (14q23.1) loci) were previously implicated across multiple studies (**Supplementary Table 4**). Over half of our regional brain sulcal associations identified were not previously implicated in any published brain imaging related studies.

### Coding variant associations

We also examined whether any of the lead variants were in strong LD (r^2^>0.8) with coding variants (*p*_discovery_<5×10^−8^ and *p*_replication_<0.05). We identified 10 loci harbouring coding variants or proxies (coding/splice region variants) in strong LD with lead variants (**Supplementary Table 5**). With the exception of the complex chr17:47Mb (17q21.31, *MAPT*) locus, which contained coding/splice region proxies for multiple genes (*ARHGAP27, PLEKHM1, CRHR1, SPPL2C, MAPT, STH, KANSL1*), the other 9 loci contained coding variants affecting proxies for single genes (*ROR1* [rs7527017, Thr518Met], *THBS3* [rs35154152, Ser279Gly], *SLC6A20* [rs17279437, Thr199Met], *EPHA3* [rs35124509, Trp924Arg], *MSH3* [rs1650697, Ile79Val], *GNA12* [rs798488, start-lost], *PDGFRL* [rs2705051, splice region variant], *EML1* [rs34198557, Ala377Val] and *TSPAN10* [rs6420484, Tyr177Cys; rs1184909254/rs10536197, frameshift indel with stop codon gained]). Notably, the *SLC6A20* Thr199Met (rs17279437) variant, associated with widespread reductions in sulcal width (**Supplementary Table 5**), has previously been associated with reduced thickness of retinal components and with increased glycine and proline derivatives in CSF and urine, consistent with the role of SLC6A20 as co-transporter regulating glycine and proline levels in the brain and kidneys, highlighting proline/glycine pathways in regulating brain sulcal widths (see **Supplementary Information** for details).

### Genetic and phenotypic correlations of brain folding

We investigated the phenotypic and genetic correlation (GC) between measures from the right and left hemispheres as well as between different shape parameters of the brain sulcal measurements. We found high correlations between brain sulcal measurements across left and right sides, within and between the four shape parameters (**Figure 1c, Extended Data Figure 7**). In general, the strongest correlations were detected between left and right hemispheres for width compared to length, mean depth and surface area (**Figure 1c top**). The high genetic correlation between hemispheres may explain the higher magnitudes of the association Z-scores of bilateral brain sulcal measures compared to hemisphere-specific analyses. We found average length, mean depth and surface area parameters to be positively correlated, with correlation between length and surface area the strongest, and width to be negatively correlated with the other 3 shape parameters (**Figure 1c middle and bottom**). Similar patterns of correlations between shape parameters were seen for left and right hemispheres as well as for both genetic and phenotypic correlations (**Figure 1c middle and bottom, Extended Data Figure 7**).

### Brain folding genes enriched for cortical expression and neurodevelopmental processes

To determine whether genes in the associated regions were enriched for expression in certain tissues, we performed enrichment analysis of annotated genes in significant loci (*p*<5×10^−8^) for tissue gene expression in an independent dataset (Human Protein Atlas) (**Methods**). We found significant enrichment of brain folding genes of approximately two-fold for expression in the cerebral cortex after multiple testing correction (*p*=7.3×10^−7^). This effect remained significant with other sensitivity analysis thresholds (**Figure 2a**), suggesting associated brain folding genes may have local effects. We also performed enrichment analysis for gene ontology (GO) processes and KEGG pathways. Notably, we found significant (FDR<0.05) enrichment for various neurodevelopmental processes including neurogenesis and a range of cellular GO biological processes: synapse, neuronal part, plasma membrane, cell junction, cytoskeletal, chromosomal and endoplasmic reticulum GO cellular components; protein domain-specific binding GO molecular function; and the axon guidance KEGG pathway (**Figure 2b**). We examined the extent and timing of expression of candidate genes across brain developmental stages using BrainSpan data in FUMA^18^ and found significant enrichment (FDR<0.05) for downregulated differentially expressed genes in early infancy (**Figure 2c**). Expression levels of numerous genes, including *DAAM1, NT5C2, NEO1* recently linked to cortical development, are downregulated during the late pre-natal 26 weeks post-conception) to early post-natal (4 months of age) period (**Extended Data Figure 8**). These results together suggest that genetic effects on regional brain folding are in part driven via regulation of neuronal development during early brain development.

**Figure 2.**
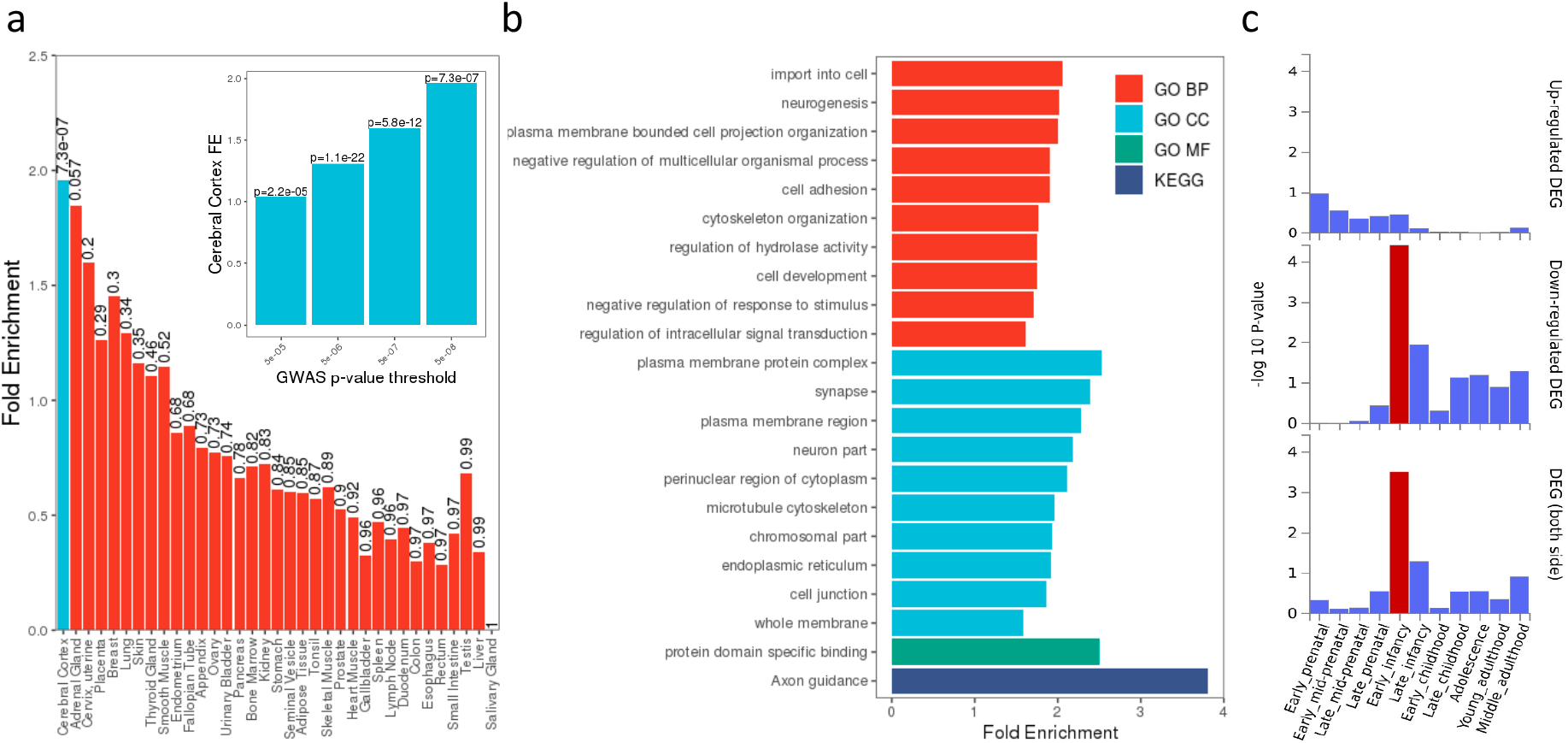
Enrichment of genes in significant loci for: **(a) gene expression across various tissues** (inset shows sensitivity analysis at other GWAS thresholds), **(b) GO and KEGG pathways** (FDR<0.05), **(c) differentially expressed genes across brain development stages**.

### Colocalization with brain eQTLs to prioritize candidate genes

We performed colocalization analysis between brain cortical folding loci and the largest cortical eQTL summary dataset generated to date (Metabrain)^19^. We found 27 of 119 loci to be colocalized for at least one sulcal measure with one or more *cis* eQTLs in the cerebral cortex at a posterior probability (coloc PP4) >0.7 and an additional 7 at a suggestive PP4 >0.5 (**Supplementary Table 6, Extended Data Figure 9**). A total of 53 unique cortical gene eQTLs colocalized (PP4 >0.7) with at least one sulcal trait in the cortex. 15 of the 27 loci were colocalized with one unique eQTL in the cortex, 9 loci colocalized with 2 eQTLs, 3 with 2 eQTLs and the pleiotropic chr17:47Mb *MAPT-KANSL1* locus colocalized with 14 different eQTL genes in a complex pattern (**Extended Data Figure 9, Supplementary Table 6**). Across other brain-related tissues including the cerebellum, basal ganglia, hippocampus and spinal cord, we found a total of 25 loci in the cerebellum, 7 in the basal ganglia, 6 in the hippocampus and 3 in the spinal cord that colocalized (PP4 >0.7) with at least one eQTL, with 9, 2 and 1 colocalized loci in the cerebellum, hippocampus and basal ganglia respectively, not found in cortex tissue.

### Multi-trait colocalization of cortex specific *KCNK2* eQTL and regional sulcal widths

The pleotropic chr1:215Mb locus near *KCNK2* is associated with multiple sulcal measures across the brain in a largely symmetrical manner. The strongest lead variant ∼40Kb upstream of *KCNK2*, rs1452628:T, exhibited stronger associations with reduced sulcal widths in more superior regions of the brain (**Figure 3a, Supplementary Tables 2 and 3**). Notably, we observed multiple pairwise colocalizations between significant sulcal width associations at this locus and cortex-specific *KCNK2* eQTLs from a large-scale brain tissue eQTL study (MetaBrain)^19^ (**Extended Data Figure 9, Supplementary Table 6**), where rs1452628:T was associated with increased *KCNK2* expression in the cortex only (beta=0.14, *p*=8.0×10^−7^) (cf. cerebellum, hippocampus, basal ganglia and spinal cord, all *p*>0.1, **Figure 3b left**). We then formally tested whether all or one or more subgroups of the regional sulcal width associations in the locus and cortical *KCNK2* eQTL are driven by the same underlying variant using the HyPrColoc multi-trait colocalization approach^20^. We found all associations multi-colocalized to the same variant (posterior probability of colocalization=0.74), with the candidate causal variant, rs1452628, explaining all of the posterior probability of colocalization (**Figure 3b right**). We further assessed sensitivity to our choice of prior probability of colocalization. Joint colocalization across all or almost all of the traits remained even after sequentially reducing the prior probability (**Supplementary Information**). These results suggest a shared underlying variant driving all sulcal morphology associations and cortex-specific *KCNK2* expression at this locus.

**Figure 3.**
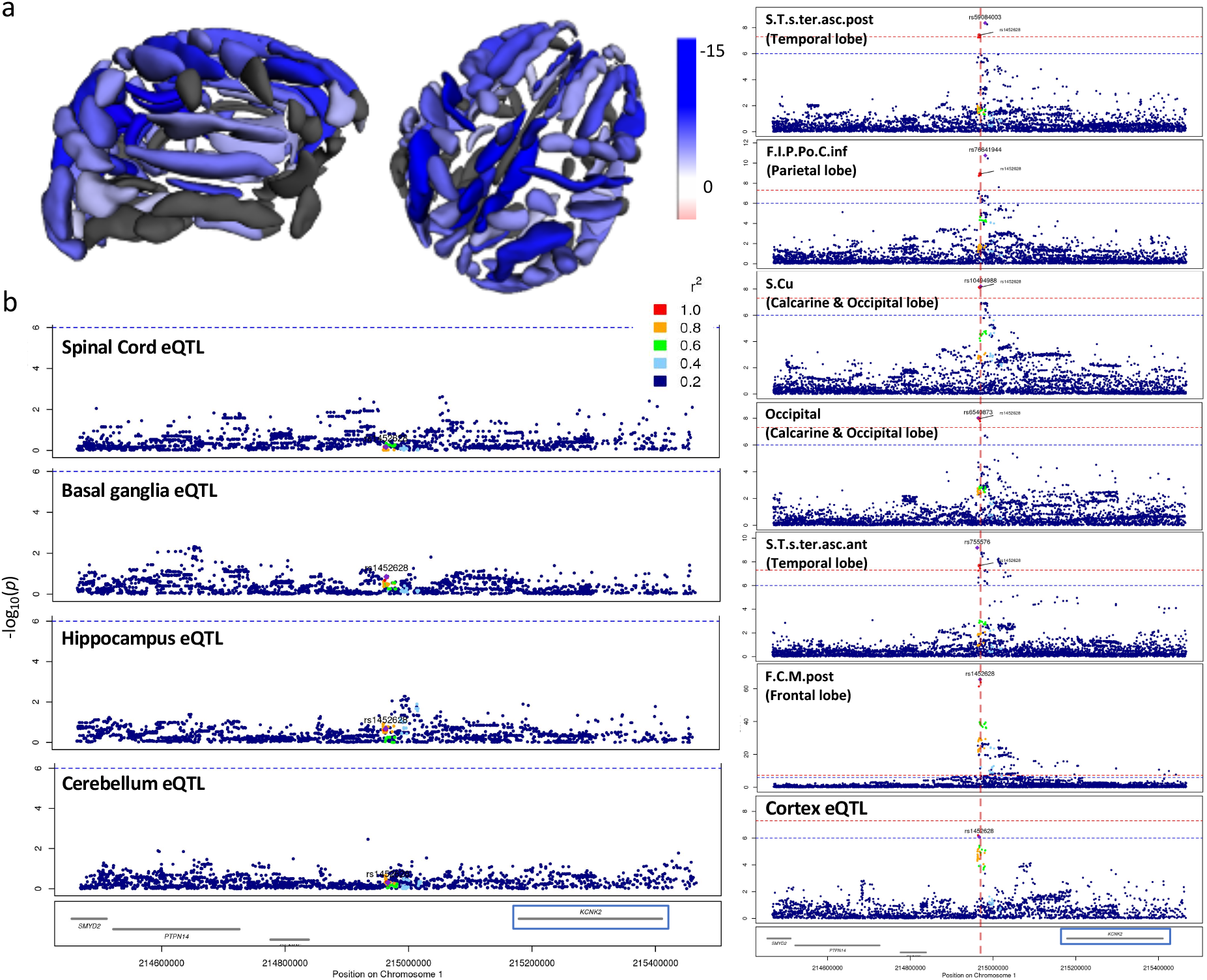
*KCNK2* locus associations. **(a) Association of the lead rs1452628:T variant with reduced sulcal widths across the brain**. (Grey colours indicate associations with *p*_*rep*_*>*0.05). **(b) Left: regional association plot of MetaBrain *KCNK2* eQTLs for spinal cord, basal ganglia, hippocampus and cerebellum. Right: regional association plots and colocalization of cortex *KCNK2* eQTL and different lead variants in the *KCNK2* locus**. A subset of associations shown for each different lead variant shown due to space constraints.

### Genetic correlation between brain folding associations and neuropsychiatric conditions

Cross referencing with previous non-imaging trait and diseases in the GWAS Catalog, we found that 56 of the 119 loci were associated with one or more diseases or intermediate phenotypes (**Supplementary Table 7**). We further investigated the genetic correlation (GC) of regional brain folding with 12 neurological diseases, cognitive and psychiatric conditions (**Methods, Supplementary Information**). Using an empirical permutation threshold of *p*<0.0044 to account for extensive correlations within brain folding phenotypes and neuro-related illnesses (**Methods**), we observed 158 significant GCs between regional brain folding measures and 10 neuropsychiatric and cognitive conditions (**Supplementary Table 8**).

Taking the mean GC between each of the four shape parameters and neuropsychiatric conditions, we found at least two distinct clusters, with generalized anxiety disorder (GAD), attention deficit hyperactive disorder (ADHD), and major depressive disorder (MDD) and Alzheimer’s disease (AD) similarly clustered (**Figure 4a**). In general, sulcal width measures mostly showed opposite GCs versus the other three sulcal parameters (**Figure 4a**), in keeping with their correlation structure. Cognitive performance and Parkinson’s disease (PD) in particular showed significant positive GCs with length, surface area and mean depth measures across a broad range of brain regions, whilst ADHD and MDD showed negative GCs across those three shape parameters (**Supplementary Table 8, Figure 4a and 4b**). In particular, we found strongest GCs between PD and central sulcal length (r_G_=0.40, *p=*3.0×10^−3^) and surface area (r_G_=0.33, *p=*6.0×10^−4^) (**Supplementary Table 8**), which indicate the role of sulcal folds adjoining the primary motor cortex in PD. Sulcal width measures mostly showed opposite GCs with neuropsychiatric traits compared the other three sulcal parameters (**Figure 4a**), in keeping with their correlation structure.

**Figure 4.**
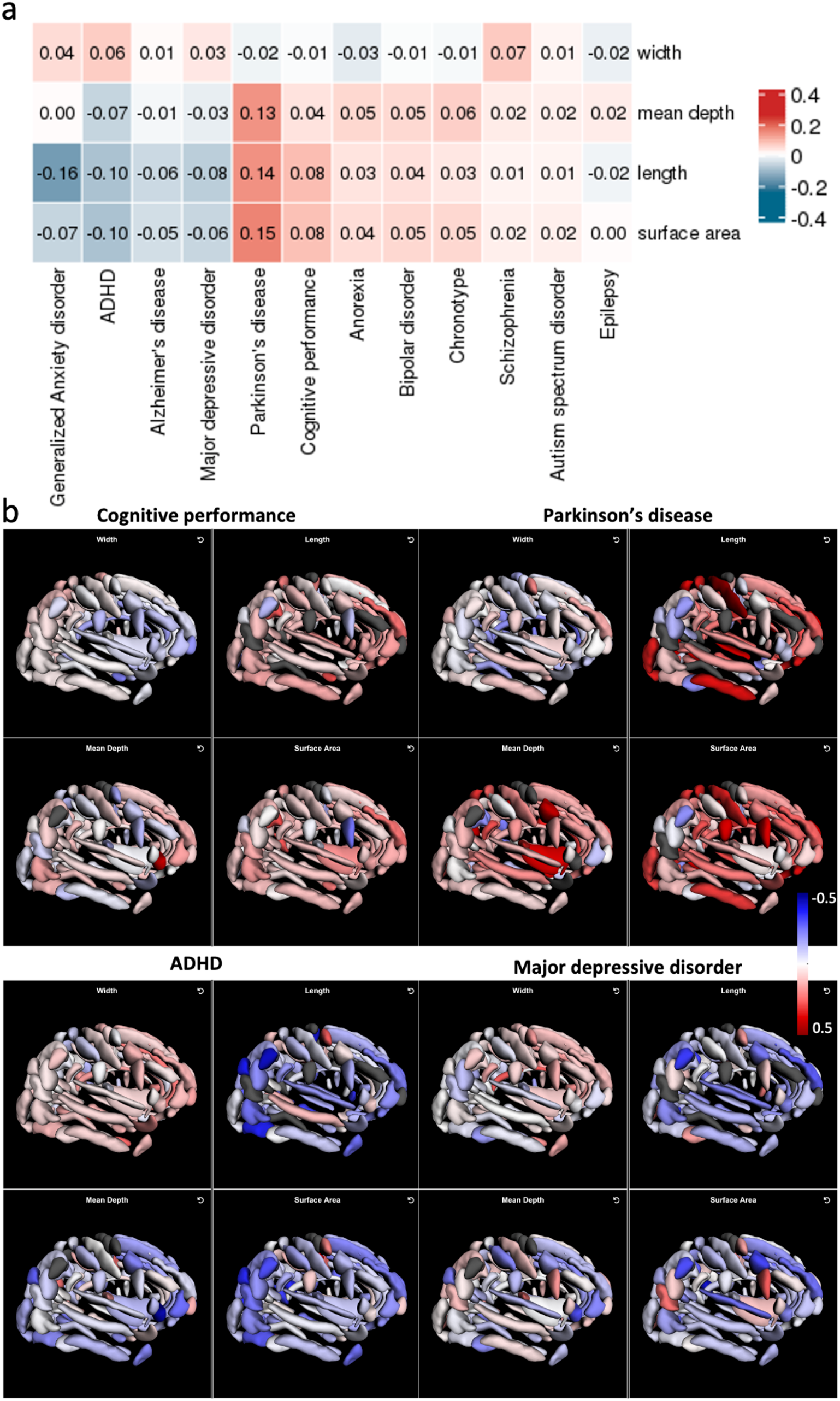
**(a) Genetic correlations between shape parameters and neuropsychiatric conditions. (b) Examples of genetic correlations across brain sulcal folds with cognitive performance, Parkinson’s disease, attention deficit hyperactive disorder (ADHD) and major depressive disorder**.

### Interactive 3D visualisation of associations

Given the complexity and interdependencies of regional brain folding, visualizing variant association results interactively in 3D provides more intuitive context to interpret the association results, providing insights into genetic effects across multiple brain regions. We created an interactive resource (https://enigma-brain.org/sulci-browser) where users can query individual genetic variants and visualize the genetic effects across all regional brain folds interactively across all four shape parameters (**Figure 5**).

**Figure 5.**
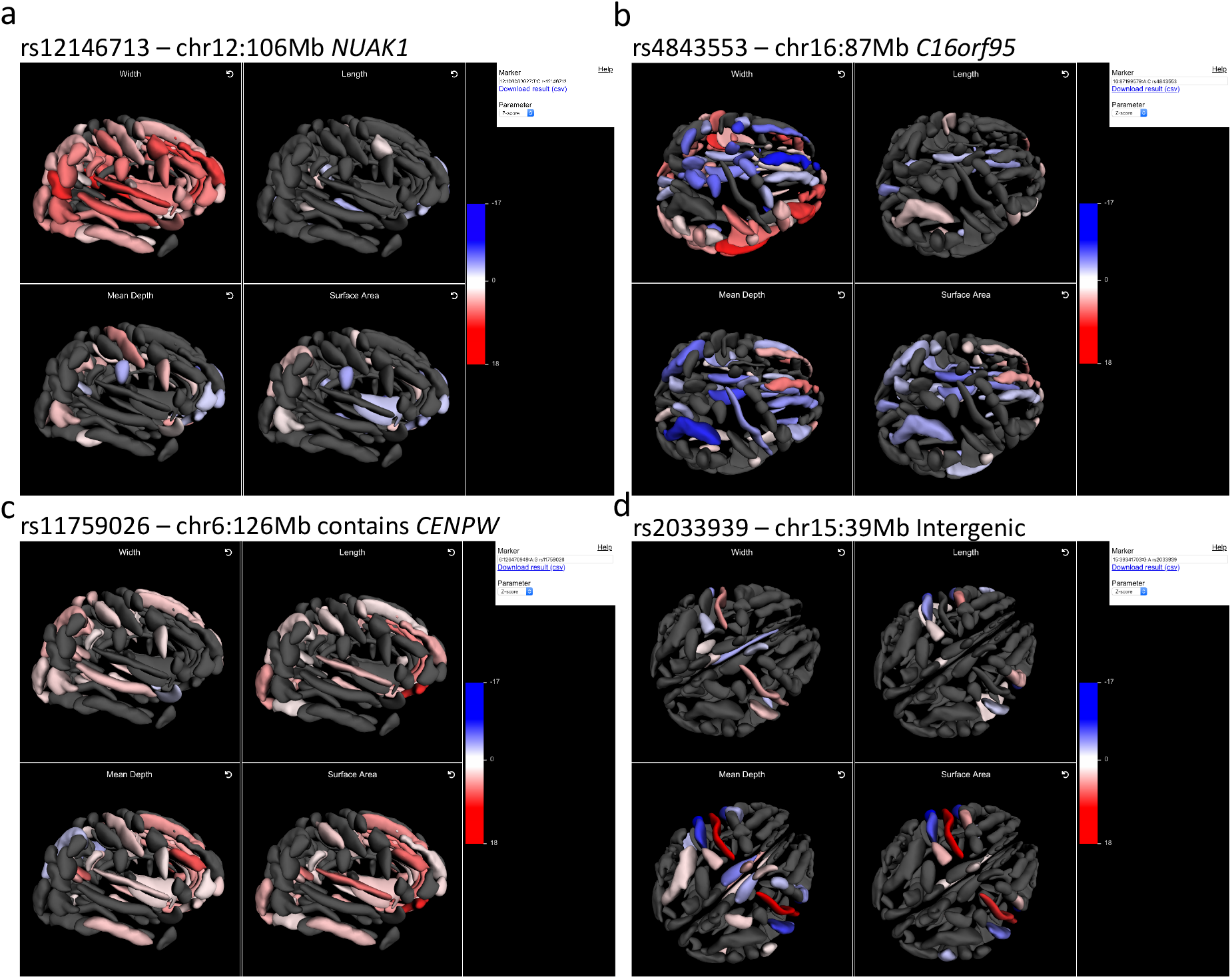
Three-dimensional visualisation of brain sulcal associations (Z-scores) for four exemplar pleiotropic loci.

Visualizing the results, for example, it is clear that pleiotropic associations, such as chr12:106Mb (*NUAK1*), chr16:87Mb (near *C16orf95*) and chr6:126Mb (containing *CENPW*) affect multiple brain regions and shape parameters in distinct and complex ways (**Figure 5a-c)**. In contrast, the chr15:40Mb (15q14) locus associations, mostly tagged by rs4924345, are more localised (**Figure 5d**). We observed strong *positive* effects of the minor rs4924345:C allele on bilateral central sulcus mean depth (beta_dis_=0.29, *p*_dis_*=*3.1×10^−79^) and surface area (beta_dis_=0.15, *p*_dis_*=*6.0×10^−25^) but *negative* effects bilaterally on neighbouring superior postcentral intraparietal superior sulcus mean depth (beta_dis_=-0.14, *p*_dis_*=*1.0×10^−18^) and surface area (beta_dis_=-0.11, *p*_dis_*=*6.5×10^−13^); retro central transverse ramus of the lateral fissure mean depth (beta_dis_=-0.16, *p*_dis_*=*7.9×10^−21^) and surface area (beta_dis_=-0.15, *p*_dis_*=*2.8×10^−18^); inferior precentral sulcus mean depth (beta_dis_=-0.14, *p*_dis_*=*5.6×10^−16^), surface area (beta_dis_=-0.16, *p*_dis_*=*8.0×10^−19^) and length (beta_dis_=-0.11, *p*_dis_*=*1.3×10^−9^).

We have also provided rendering based on effect sizes, Z-scores or *p*-values and an option to download query results.

## Discussion

Cortical gyrification is an orchestrated, multifaceted process that shows striking consistency across individuals^21^. Gyrification is regulated by a complex interplay of cellular, biomechanical and genetic influences^9^ but our understanding of its genetic underpinnings has been limited^22,23^. Abnormalities and variations in brain folding contribute to many common and rare neuropsychiatric conditions. Cortical thickness, surface area and sulcal morphometry are each associated with complex phenotypes such as intelligence^24^, and effects on cortical gyrification are partially independent of those on cortical thickness or surface area^25^.

Here, combining densely-imputed genetic variants with whole-exome sequencing, we performed the most comprehensive genetic mapping of regional cortical sulcal morphometry to date, identifying 119 unique genetic loci influencing human sulcal depth, width, length and surface area. We discovered over 60 novel loci not previously implicated in any brain imaging related association studies. The number of genetic associations observed across different sulcal parameters was approximately in accordance with their heritability^17^. We observed stronger genetic correlations than phenotypic correlations between left and right sides, suggesting that environmental and non-genetic factors may play a role in structural and functional lateralization. In particular, regional measures for the most heritable shape parameter, sulcal width, clustered in a way that reflected broad brain topology, indicating that brain sulcal width has a stronger genetic component and is most stable across the lifespan.

We demonstrated the highly polygenic genetic architecture of brain folding, which has both local and widespread effects within the brain. When visualised in 3D, local effects are apparent, that are likely to be missed in globally aggregated brain measurement studies. We also implicated specific candidate genes in several cases through coding variants in LD. We added exonic resolution through WES, as well as through colocalization with brain eQTLs using a large-scale brain specific dataset for better power and specificity^19^. We observed pleiotropic associations at genetic loci consistently implicated in prior genetic studies of neuroimaging phenotypes, such as the *MAPT-KANSL1* locus^26,27^, while resolving other associations to specific brain regions and sulcal folding parameters, such as the *KCNK2* locus and sulcal width.

Our results provide evidence of enrichment of associated genes for expression in the cerebral cortex, strongly implicating genes involved in neurodevelopment. We found enrichment for differential gene expression occurring in early brain development, indicating that genetic effects on cortical gyrification occur most prominently during early life, likely via modulation of neurodevelopmental pathways. Inherited functional impairments of these genes and their associated pathways may increase the risk for neurodevelopmental disorders. For example, homozygous and compound heterozygous mutations at *EML1 -* a gene associated with right insula surface area - cause band heterotopia, a neuronal migration disorder characterized by intellectual disability and epilepsy^28^. Similarly, heterozygous deletion of *ZIC1* and *ZIC4* is associated with Dandy-Walker malformation, a congenital cerebellar malformation^29^, whereas contiguous deletions at the 16q24.3 locus encompassing *CENPW* cause microcephaly, distichiasis, vesico-ureteral and intellectual impairment^30^. Additionally, genetic variants at *NUAK1 -* a pleiotropic locus associated with frontal, temporal and precentral sulcal widths - have shown links to autism spectrum disorder^31,32^, ADHD^33^ and cognitive impairment^34^. Globally, genetic variants influencing cortical gyrification showed robust, widespread correlation with variants influencing cognitive performance, schizophrenia, ADHD and depression, suggesting a shared molecular system potentially underpinning neurodevelopmental and neuropsychiatric disorders^35,36^.

Through multi-trait colocalization, we identified a shared underlying genetic driver of increased cortical *KCNK2* expression and pleiotropic effects on reduced sulcal widths. KCNK2, also known as TREK-1, is a two-pore domain potassium channel highly expressed in the central nervous system and modulated by both chemical and physical stimuli.^37,38^ KCNK2 regulates immune-cell trafficking into the CNS^37^ and genetic ablation of *Kcnk2* is associated with neuroinflammation, blood-brain barrier impairment^39^ and increased sensitivity to ischemia and epilepsy in mice^40^. In addition to brain volume, the *KCNK2* locus was previously implicated in sulcal opening^16^ and the same lead variant, rs1452628:T, was associated with difference between predicted brain age and chronological age^41^. Our findings re-emphasize the role of *KCNK2* in cerebral cortex development, alongside similarly pleiotropic and widely-investigated therapeutic targets such as *NUAK1*^42^ and *MAPT*^*43*^. Further investigation of the links between these proteins and disease processes downstream of cortical gyrification may support therapeutic development.

One notable limitation of the present study is that genetic associations were identified in a population of mostly British individuals. Additionally, dividing UK Biobank participants into discovery and replication cohorts prioritised robustness of genetic associations, but reduced power to detect rare and low frequency variant associations. Larger sample sizes will increase power and refine the estimates reported here. Our method to ascertain brain folding phenotypes is applicable across different MRI scanning protocols, which vary across sites^17^. This should facilitate large-scale, cross-biobank studies of cortical folding and minimise site- and cohort-specific effects.

To aid interpretation and increase the utility of our results to the wider scientific community, we created an interactive 3D brain visualisation of our associations, where users can query specific variant associations across the entire brain and the shape parameters simultaneously. We highlighted various cases where complex and pleiotropic associations differ in brain region and shape parameter distributions, which become more apparent when represented visually in three dimensions.

In conclusion, we provide the most comprehensive genetic atlas of regional brain folding to date, identifying novel associations and insights into processes that drive the genetic effects, as well as providing a resource for the wider community for further elucidation of specific findings.

## Methods

### Samples and participants

UK Biobank (UKB) is a UK population study of approximately 500,000 participants aged 40-69 years at recruitment^44^. Participant data include genomic, imaging data, electronic health record linkage, biomarkers, physical and anthropometric measurements. Further details are available at https://biobank.ndph.ox.ac.uk/showcase/. Informed consent were obtained from participants. Analyses in this study were conducted under UK Biobank Approved Project numbers 26041 and 11559.

### Brain folding imaging phenotypes

The UK Biobank began collecting brain MRI scans in 2014 with the goal of scanning 100,000 individuals. The protocol includes isotropic 3D T1-weighted (T1w) MP-RAGE images (voxel size 1 mm^3^; field-of-view: 208 × 256 × 256) that have undergone bias-field correction in the scanner. Full acquisition details can be found in^45^. T1w images were processed using FreeSurfer (v7.1.1) (https://surfer.nmr.mgh.harvard.edu/) and quality controlled using protocols developed by the Enhancing Neuro Imaging Genetics for Meta-Analysis (ENIGMA) consortium (http://enigma.ini.usc.edu/). BrainVISA (http://brainvisa.info) was implemented for sulcal classification and labelling^46^. Morphologist 2015, an image-processing pipeline included in BrainVISA, was used to measure sulcal shape descriptors. To improve sulcal extraction and build on current protocols used to analyse thousands of brain scans, quality controlled FreeSurfer outputs (*orig*.*mgz, ribbon*.*mgz and talairach*.*auto*) were directly imported into the pipeline to avoid re-computing intensities inhomogeneities correction and grey/white matter classification. Sulci were then automatically labelled according to a predefined anatomical nomenclature^46,47^. This protocol is part of the ENIGMA-SULCI working group; a Docker and a Singularity container have been created to facilitate the processing on computational clusters (https://hub.docker.com/repository/docker/fpizzaga/sulci). We retained length, width, depth, and surface area for all 121 sulcal measurements derived from this protocol for a total of 484 phenotypes.

Phenotypes with missingness >75% were excluded from subsequent analysis, leaving 450 measurements for analysis. Missingness occurs mainly with smaller sulci that are not identified in some individual MRIs. Prior to analysis, all imaging phenotypes were inverse-rank normalised to approximate a standard normal distribution and minimise effects of outliers. T-distributed stochastic neighbour embedding (t-SNE) was applied on inverse-rank normalised imaging phenotypes.

### Discovery and replication cohorts

We partitioned UKB samples with MRI measurements into discovery and replication approximately in 2:1 split. The discovery cohort were comprised of MRI measures in individuals of European ancestry from Newcastle, Stockport and Reading imaging centres, whilst the replication cohort composed of the remaining (non-European) individuals from the aforementioned three centres, and mostly all individuals from the Bristol imaging centre. Subsequent analyses were performed treating the discovery and replication cohorts as completely separate to minimize data contamination and biases.

### Genetic data processing

#### UKB genetic QC

UKB genotyping and imputation (and QC) were performed as described previously^44^. WES data for UKB participants were generated at the Regeneron Genetics Center (RGC) as part of a collaboration between AbbVie, Alnylam Pharmaceuticals, AstraZeneca, Biogen, Bristol-Myers Squibb, Pfizer, Regeneron and Takeda with the UK Biobank^48^. WES data were processed using the RGC SBP pipeline as described in^49,50^. RGC generated a QC-passing “Goldilocks” set of genetic variants from a total of 454,803 sequenced UK Biobank participants for analysis. Additional QC were performed prior to association analyses as detailed below.

#### Additional QC and variant processing

In addition to checking for sex mismatch, sex chromosome aneuploidy, and heterozygosity checks, imputed genetic variants were filtered for INFO>0.8, MAF>0.01 (rarer variants around coding regions would be better captured by WES) globally across UKB and chromosome positions were lifted to hg38 build. WES variants were filtered for MAC>10 within the UKB subset with MRI measurements. Imputed and WES variants were combined by chromosome position (hg38) and alleles and in the case of overlaps, the WES variant was retained (as WES generally have higher quality calls compared to imputation). Variant annotation was performed using VEP^51^ with Ensembl canonical transcripts used where possible.

### Genetic association analyses

GWAS were performed using REGENIE v2.0.1 via a two-step procedure to account for population structure detailed in^52^. In brief, the first step fits a whole genome regression model for individual trait predictions based on genetic data using the leave one chromosome out (LOCO) scheme. We used a set of high-quality genotyped variants: minor allele frequency (MAF)>1%, minor allele count (MAC)>100, genotyping rate >99%, Hardy-Weinberg equilibrium (HWE) test *p*>10^−15^, <10% missingness and linkage-disequilibrium (LD) pruning (1000 variant windows, 100 sliding windows and r^2^<0.8). The LOCO phenotypic predictions were used as offsets in step 2 which performs variant association analyses using standard linear regression. We limited analyses to variants with MAC>50 to minimise spurious associations. The association models in both steps also included the following covariates: age, age^2^, sex, age*sex, age^2^*sex, imaging centre, intracranial volume, first 10 genetic principal components (PCs) derived from the high-quality genotyped variants (described above) and additionally first 20 PCs derived from high-quality rare WES variants (MAF<1%, MAC>5, genotyping rate >99%, HWE test *p*>10^−15^, <10% missingness) as additional control for fine-scale population structure.

### Definition and refinement of significant loci

To define significance, we used multiple testing corrected threshold of *p*<2×10^−10^ (5×10^−8^/273 approximate number of independent trait). We used phenotypic PCs accounting for 90% of phenotype variance to estimate the approximate number of independent traits to account for correlations between regions, side and parameters. Additionally, we also require at least nominal significance (*p*<0.05) with concordant directions in the replication cohort which should limit false positives even at *p<*5×10^−8^. For reporting, we also included the standard genome-wide significant loci (*p<*5×10^−8^) that replicated at *p*<0.05 in the replication cohort.

We defined independent trait associations through clumping ±500Kb around the lead variants using PLINK^53^, excluding the HLA region (chr6:25.5-34.0Mb) which is treated as one locus due to complex and extensive LD patterns. As overlapping genetic regions may be associated with multiple correlated measurements and to avoid over-reporting genetic loci, we merged overlapping independent genetic regions (±500Kb) across traits and collapsed them into one locus.

### Cross reference with known genetic associations

We cross-referenced the lead variants and their proxies (LD proxy r^2^>0.8, +/-500Kb around the lead variant, with HLA region treated as one region) for significant associations (*p*<5×10^− 8^) in GWAS Catalog^6^. Brain imaging studies were separated from other intermediate and disease phenotypes as defined by the list of brain imaging studies in **Supplementary Information**.

### Expression enrichment

We examined whether genes within associated loci are enriched for expression the various brain tissues. Enrichment analysis was performed using the TissueEnrich R package^54^ using the annotated genes (available canonical genes mapped in VEP) for all genome-wide significant variants (*p*<5×10^−8^, additional sensitivity analysis thresholds of *p*<5×10^−7^, 5×10^−6^, 5×10^−5^ were used for cortex) and a background of annotated genes for all variants analysed. Specifically, we used the RNA dataset from Human Protein Atlas using all genes that are found to be expressed within each tissue.

### GO and KEGG process enrichment

Using the same significant annotated genes and backgrounds as for the expression enrichment analyses, we performed enrichment testing for GO and KEGG pathways using the WEB-based GEne SeT AnaLysis Toolkit (WebGestalt)^55^ (http://www.webgestalt.org/). We used the over-representation analysis method, analysing GO Biological Process, GO Cellular Component, GO Molecular Function and KEGG, with Benjamini-Hochberg FDR threshold of 0.05 for significance. We used the default parameters of minimum of 5 and maximum 2,000 genes per category. Related process and pathway entries were grouped through the inbuilt weighted set cover redundancy reduction approach.

### FUMA analyses of expression timing

Gene expression enrichment across BrainSpan (https://www.brainspan.org/) brain ages and developmental stages was analysed based on averaged log2 transformed expression levels across each label. Genes were defined as differentially expressed when the Bonferroni corrected *p*<0.05 and the absolute log fold change ≥0.58 between specific brain ages or developmental stages compared to others^18^. All other annotated genes/transcripts in the BrainSpan data were included as background genes for comparison in hypergeometric tests of gene sets. Significantly enriched gene sets had FDR corrected *p*<0.05.

### Genetic correlation analysis

We performed genetic correlation analysis between brain folding phenotypes (including hemispheres and shape parameters), and 12 neuropsychiatric conditions with readily available summary data using LD score regression (LDSC v1.0.1)^56^. We also performed SNP-based heritability estimation using LDSC. Genetic variants were filtered and processed using the “munge_sumstats.py” in LDSC and we used LD scores recommended by the software authors^56^.

To account for multiple testing of extensive related and correlated phenotypes, we permuted each neuropsychiatric condition Z-score 100 times (limited by computational cost) and tested each permuted neuropsychiatric condition with each brain folding phenotype to generate an empirical multiple testing threshold of *p*=0.0044 (approximately adjusted *p*<0.01 from 100 permutations).

### Colocalization analyses

We performed colocalization analyses^57^ between brain eQTLs from MetaBrain and brain folding loci using the coloc R package. We used the default priors (p1=10^−4^, p2=10^−4^, p12=10^− 5^) with regions defined as +/-500Kb around the lead variant. Evidence for colocalization was assessed using the posterior probability (PP) for hypothesis 4 (PP4; an association for both traits driven by the same causal variant). PP4>0.5 were deemed likely to colocalize as it guaranteed that hypothesis 4 was computed to have the highest posterior probability, and PP4>0.7 were deemed highly likely to colocalize.

To assess whether all traits jointly colocalize at the *KCNK2* locus we used the multi-trait colocalization software HyPrColoc^20^, using the recommended default settings and priors (HyPrColoc’s default prior parameters p=10^−4^ and p_c_=2×10^−2^ are equivalent to setting p1=10^−4^, p2=10^−4^, p12=2×10^−6^ in coloc, hence the default prior probability of colocalization p12 is slightly more conservative than in coloc). HyPrColoc computes evidence supporting one or more clusters of traits colocalizing at a single variant in the region, concluding that a cluster of traits colocalize if the posterior probability of colocalization (PPC) is above a user defined threshold (PPC>0.5 by default, which is equivalent to setting the algorithms’ regional, P_R_, and alignment, P_A_, thresholds to 0.7 respectively). We also performed additional sensitivity analysis across different parameter specifications (**Supplementary Information**).

## Acknowledgements

We thank all the participants, contributors and researchers of UK Biobank for making data available for this study. We thank the UK Biobank Exome Sequencing Consortium (AbbVie, Alnylam Pharmaceuticals, AstraZeneca, Biogen, Bristol-Myers Squibb, Pfizer, Regeneron and Takeda) for generation the whole exome sequencing data and Regeneron Genetics Centre for initial quality control of the exome sequencing data. We thank Peter Kochunov and his team for hosting the interactive browser, with support from NIH instrumentation grant S10OD023696. N.J. is supported by NIH grant R01AG059874. S.E.M. and J.N.P. are supported in part by NHMRC grants APP1172917 and APP1158127.

## Data availability

The online browser for visualisation of results is available at https://enigma-brain.org/sulci-browser.

## Code availability

Codes used are part of standard software and tools. Additional details available in **Methods**.

## Author contributions

Study conceptualization and design: C.D.W., P.M.T., N.J., B.B.S.; methodology: B.B.S., S.J.L., P.M.T., N.J., C.D.W.; sulcal imaging processing: F.P., A.Z., D.D., T.I., I.B.G., N.J.; phenotype harmonisation: M.J., D.G.M., S.S.C., Biogen Biobank Team; analysis: B.B.S., S.J.L., J.N.P., S.E.M., C.N.F.; interactive browser: N.S., F.P.; writing: B.B.S., C.D.W., P.M.T., N.J., H.R.; all authors critically reviewed the manuscript.

## Competing interests

The authors declare the following competing interests: B.B.S., S.J.L., Biogen Biobank Team, M.J., D.G.M., H.R., C.D.W. are employees of Biogen. P.M.T and N.J received grant support from Biogen for this work.

## Notes

https://enigma-brain.org/sulci-browser

