## Supplementary Figures for "Genetic map of regional sulcal morphology in the human brain"

### Extended Data Figures

**Extended Data Figure 1. t-SNE representation of brain sulci folds coloured by shape parameters.**

**Extended Data Figure 2. GWAS Study overview.**

**Extended Data Figure 3. Manhattan plots by brain region, shape parameter and side.**

**Extended Data Figure 4. Effect size vs minor allele frequency (MAF) of sentinel associations across sides and shape parameters.**

**Extended Data Figure 5. SNP-based heritability estimates.**

**Extended Data Figure 6. Comparison of Z-scores between sides.**

**Extended Data Figure 7. Genetic and phenotypic correlation heatmap of local sulcal measures.**

**Extended Data Figure 8. Heatmap of gene expression across brain developmental stages.**

**Extended Data Figure 9. Colocalization heatmap of brain tissue *cis* eQTLs against significant brain sulcal associations.**

**Extended Data Figure 1. t-SNE representation of brain sulci folds coloured by shape parameters.**

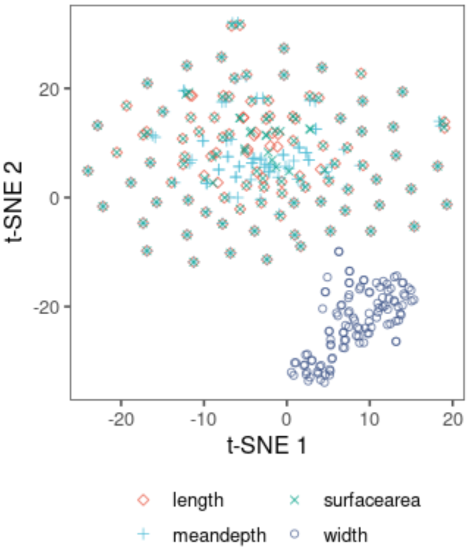

**Extended Data Figure 2. GWAS Study overview.**

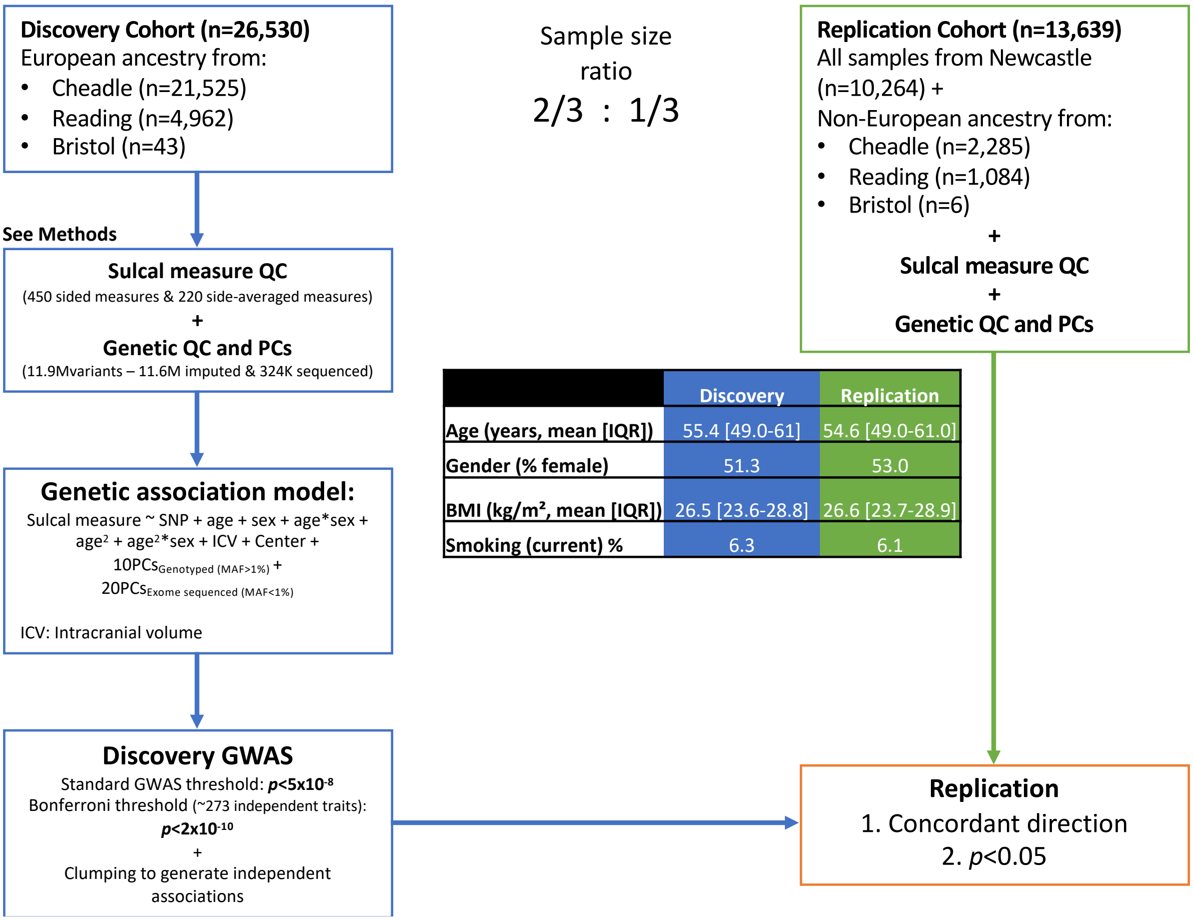


| | Discovery | Replication |
|---|---|---|
| Age (years, mean [IQR]) | 55.4 [49.0-61] | 54.6 [49.0-61.0] |
| Gender (% female) | 51.3 | 53.0 |
| BMI (kg/m², mean [IQR]) | 26.5 [23.6-28.8] | 26.6 [23.7-28.9] |
| Smoking (current) % | 6.3 | 6.1 |

**Extended Data Figure 3. Manhattan plots by brain region, shape parameter and side.** Points above 0 in the y-axis in each plot refers to associations with **left** sided sulcal measures, below 0 with **right** sided measures. Diamonds along 0 is the y-axis indicate sentinel associations for **bilateral** sulcal measures.

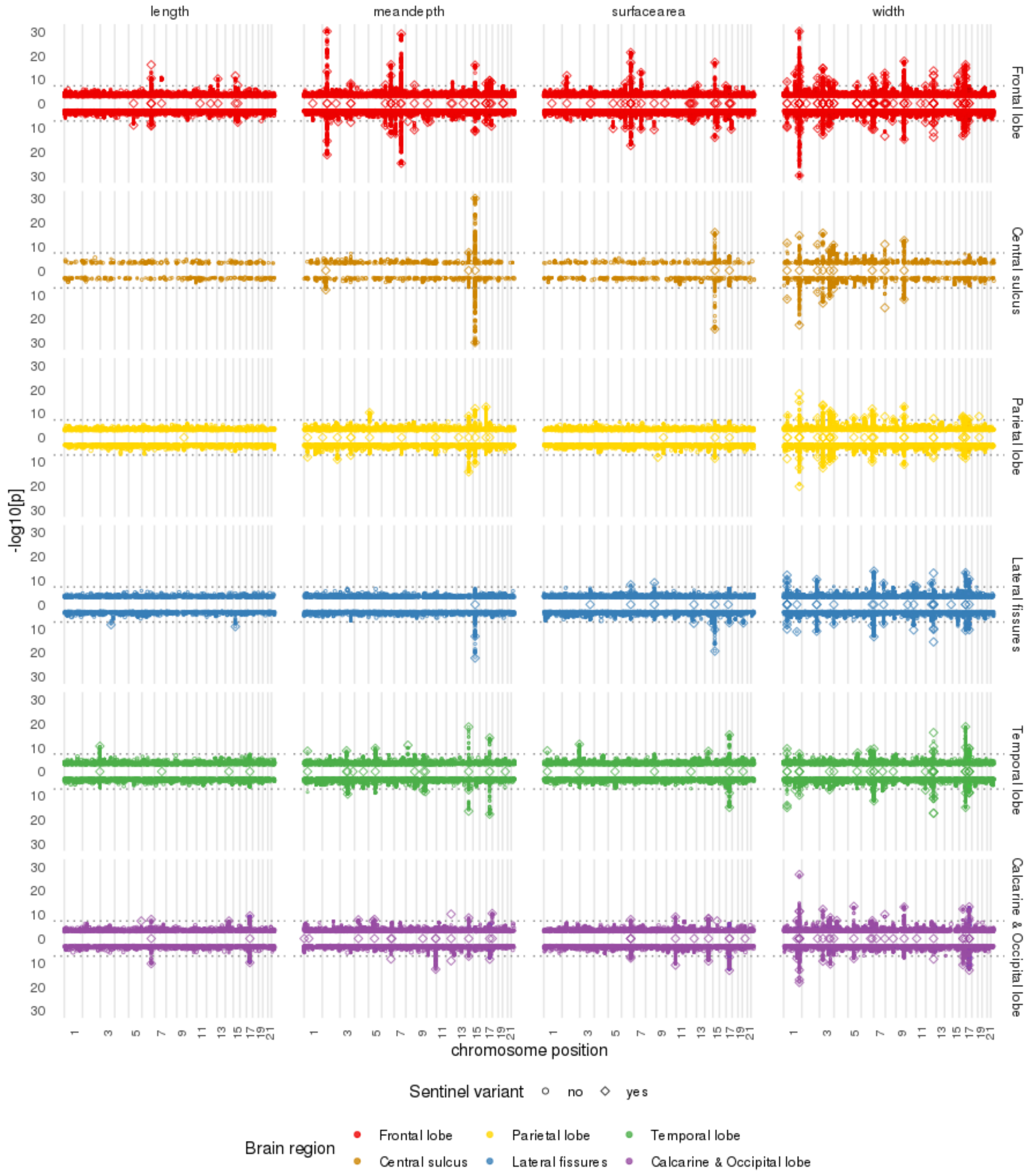

**Extended Data Figure 4. Effect size vs minor allele frequency (MAF) of sentinel associations across sides and shape parameters.**

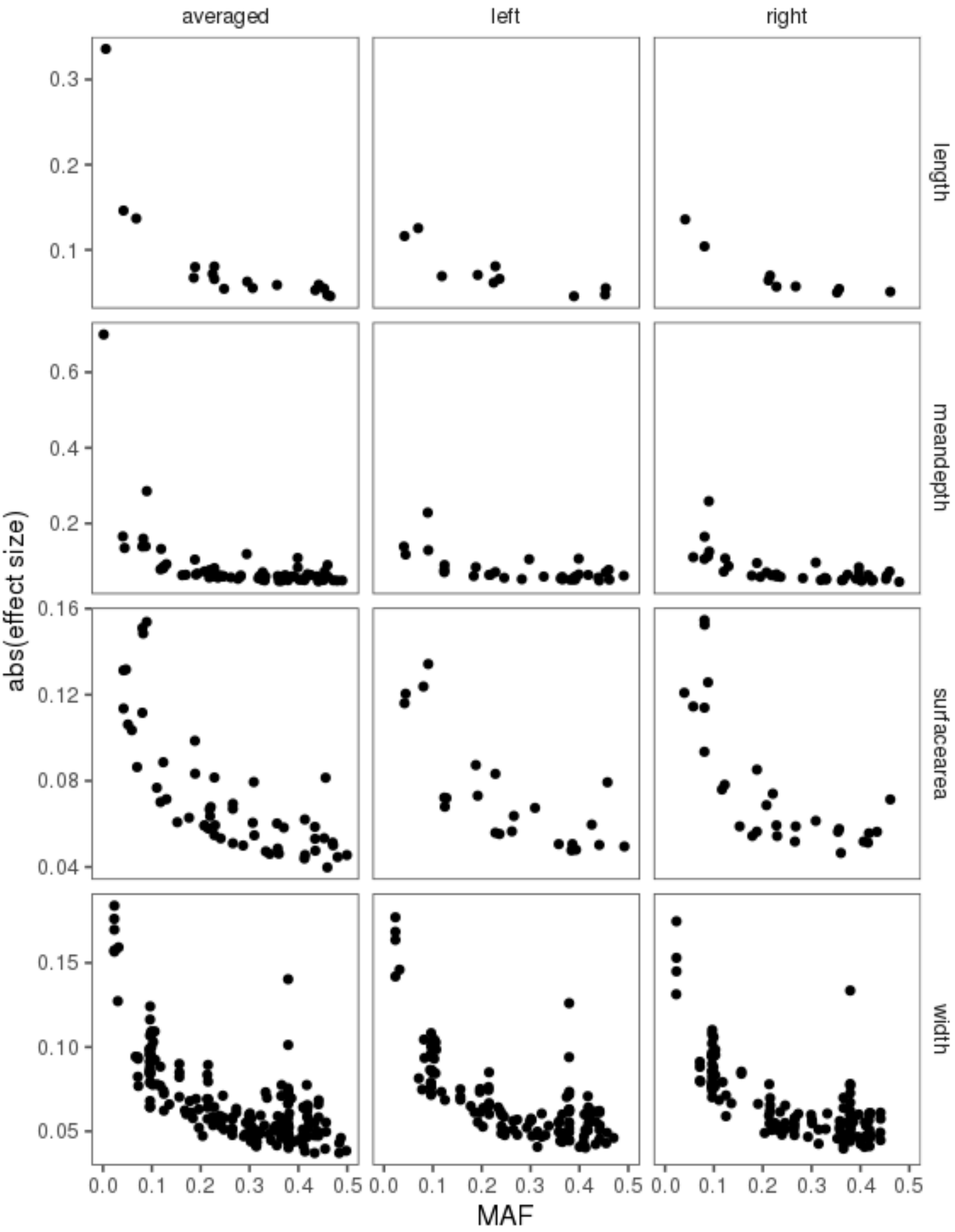

**Extended Data Figure 5. SNP-based heritability estimates.**

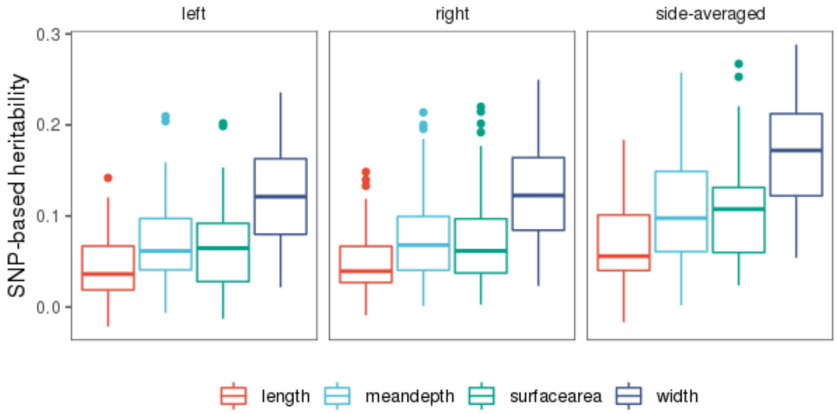

**Extended Data Figure 6. Comparison of Z-scores between sides.** Z_left, Z_right, Z_mean correspond to Z-scores of left, right and bilateral sulcal measures respectively.

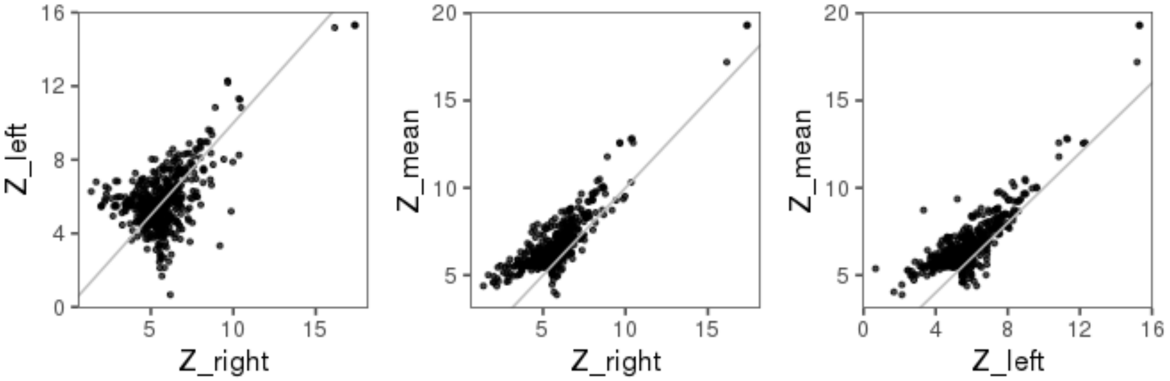

**Extended Data Figure 7. Genetic** (top triangle) **and phenotypic** (bottom triangle) **correlation heatmap of local sulcal measures.**

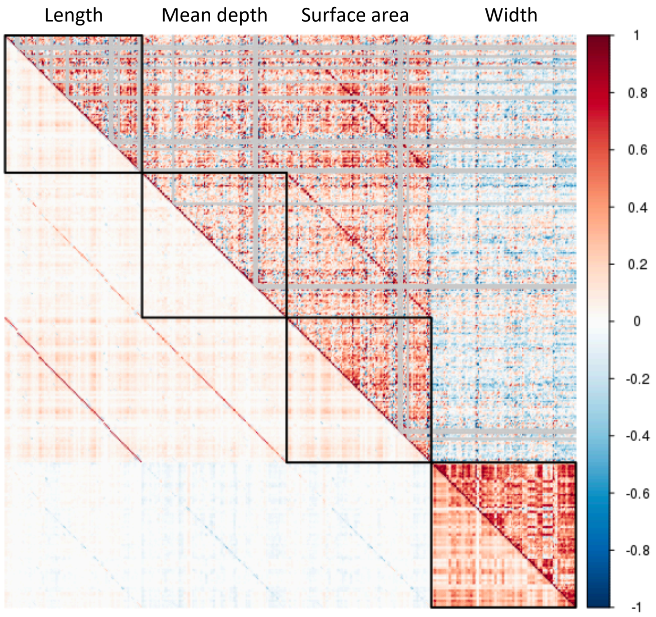

**Extended Data Figure 8. Heatmap of gene expression across brain developmental stages**. Log2 transformed BrainSpan expression values (RPKM; Read Per Kilobase per Million) for all genes in significant loci.

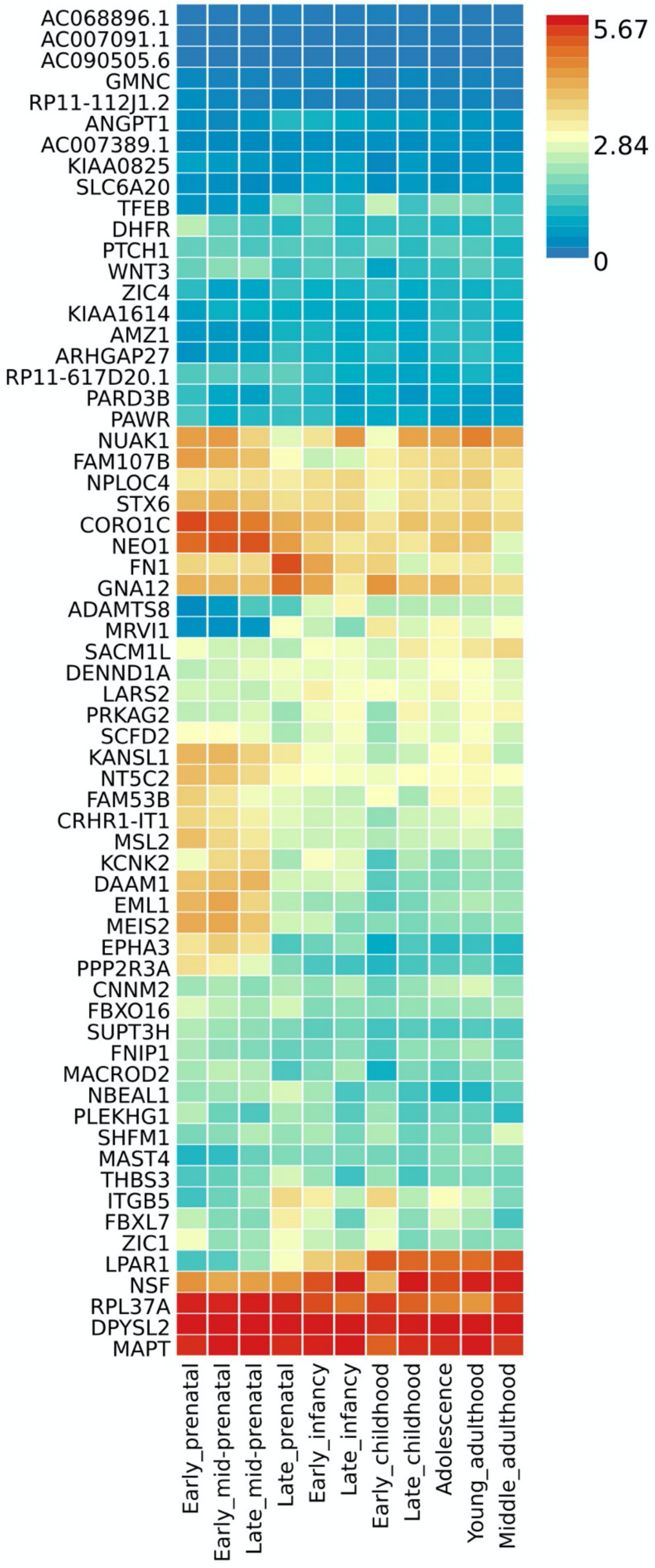

**Extended Data Figure 9. Colocalization heatmap** (PP4>0.5 shown) **of brain tissue *cis* eQTLs against significant brain sulcal associations.**

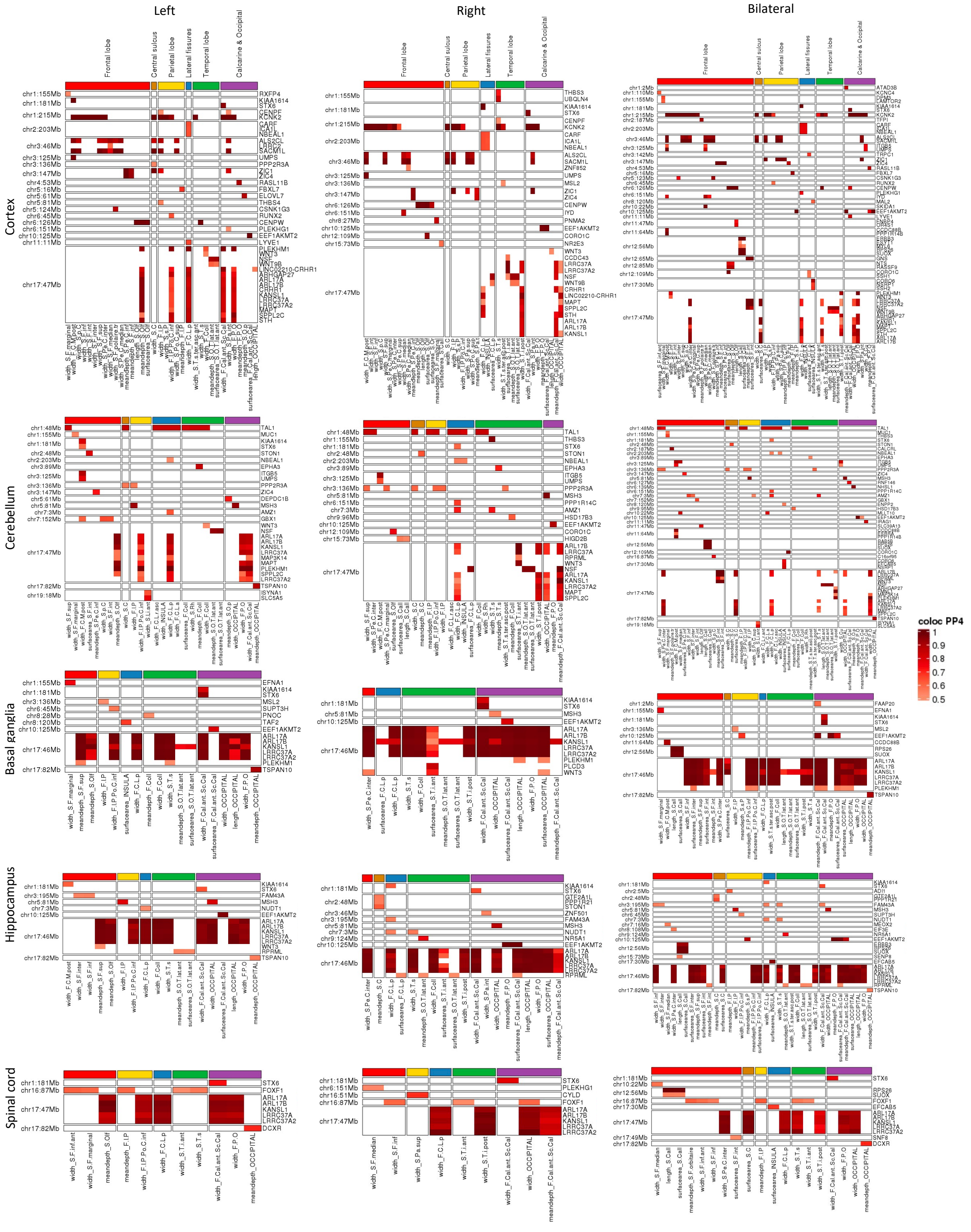
