## Supplementary Information for "Genetic map of regional sulcal morphology in the human brain"

#### Missense variant in ***SLC6A20*** transporter highlights role of glycine and proline pathways in brain sulcal width modulation

We discovered the missense variant rs17279437, causing Thr199Met in *SLC6A20*, to be associated, either as the lead variant or strong proxies ($r^2>0.8$), with widespread reduced sulcal widths (**Supplementary Table 5 and Supplementary Figure *SLC6A20***). SLC6A20 is an amino-acid (especially proline/glycine) co-transporter expressed in the kidneys, intestines and brain[1]. This transporter has recently identified to play a role in brain glycine homeostasis and NMDA-type glutamate receptor (NMDAR) function[2]. Thr199Met SLC6A20 has a greatly reduced transport capacity compared with the wild-type protein[3] and has been implicated in iminoglycinuria and hyperglycinuria (abnormally proline, hydroxyproline and glycine levels in the urine)[4], consistent with *SLC6A20* expression and reduced reabsorption in renal tubules. Metabolomic QTL (mQTL) studies have also shown *SLC6A20* Thr199Met associations with urinary levels and ratios involving glycine derivatives[5,6] and with blood levels of pyroglutamate[7]. Within the nervous system, *SLC6A20* Thr199Met has been recently associated with increased levels of betaine (trimethylglycine)[8] and L-proline[9] in cerebrospinal fluid. In addition to widespread reduced sulcal widths, the Thr199Met variant is also associated with reduced macular thickness[10], reduced retinal nerve fibre layer and ganglion cell inner plexiform layer thicknesses[11] and increased risk of macular telangiectasia type 2[12] in the eye.

Together, these results suggest the role of SLC6A20 and glycine/proline related pathways in conditions related to retinal thicknesses or reduced sulcal widths, such as macular telangiectasia (type 2) or neurodegenerative and psychiatric conditions, in addition to renal conditions such as iminoglycinuria/hyperglycinuria.

**Supplementary Figure *SLC6A20*. Association (Z-scores) of rs17279437 on sulcal widths in the brain.**

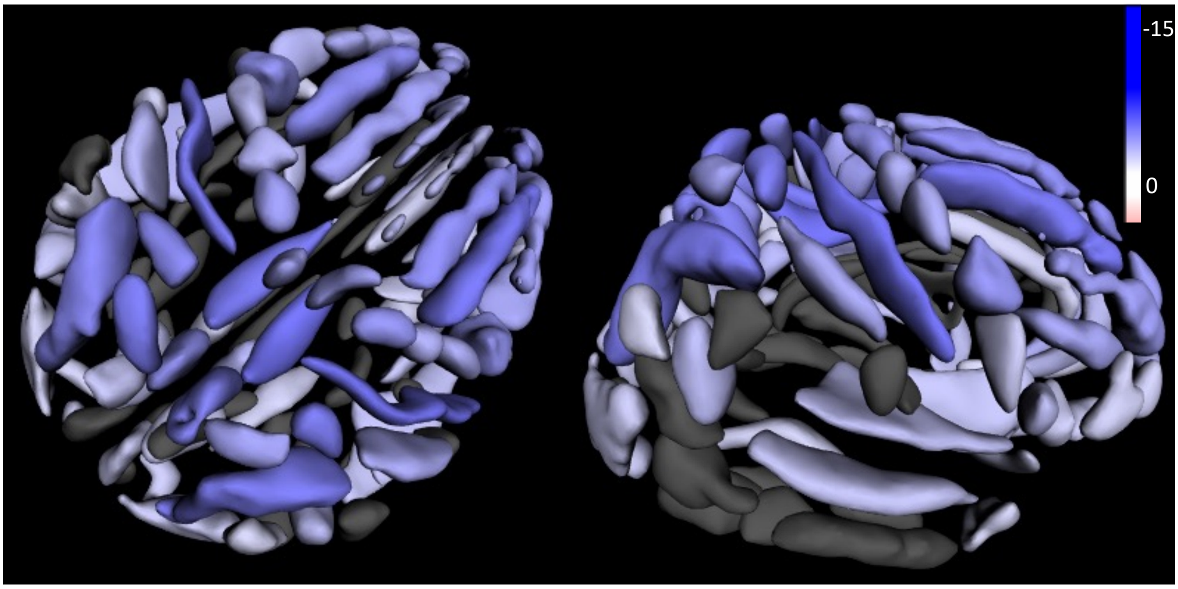

### Summary of brain imaging related studies in GWAS Catalog

| PMID | First Author | Year | Title |
|---|---|---|---|
| **33723403** | Sha Z | 2021 | The genetic architecture of structural left-right asymmetry of the human brain. |
| **33293549** | Sargurupremraj M | 2020 | Cerebral small vessel disease genomics and its implications across the lifespan. |
| **32963231** | Hofer E | 2020 | Genetic correlations and genome-wide associations of cortical structure in general population samples of 22,824 adults. |
| **32665545** | van der Meer D | 2020 | Understanding the genetic determinants of the brain with MOSTest. |
| **32358547** | Persyn E | 2020 | Genome-wide association study of MRI markers of cerebral small vessel disease in 42,310 participants. |
| **32198502** | Shin J | 2020 | Global and Regional Development of the Human Cerebral Cortex: Molecular Architecture and Occupational Aptitudes. |
| **32193296** | Grasby KL | 2020 | The genetic architecture of the human cerebral cortex. |
| **31676860** | Zhao B | 2019 | Genome-wide association analysis of 19,629 individuals identifies variants influencing regional brain volumes and refines their genetic co-architecture with cognitive and mental health traits. |
| **31666681** | Zhao B | 2019 | Large-scale GWAS reveals genetic architecture of brain white matter microstructure and genetic overlap with cognitive and mental health traits (n=17,706). |
| **31636452** | Satizabal CL | 2019 | Genetic architecture of subcortical brain structures in 38,851 individuals. |
| **31396565** | van der Lee SJ | 2019 | A genome-wide association study identifies genetic loci associated with specific lobar brain volumes. |
| **30818988** | Klein M | 2019 | Genetic Markers of ADHD-Related Variations in Intracranial Volume. |
| **30649180** | Luo Q | 2019 | Association of a Schizophrenia-Risk Nonsynonymous Variant With Putamen Volume in Adolescents: A Voxelwise and Genome-Wide Association Study. |
| **30305740** | Elliott LT | 2018 | Genome-wide association studies of brain imaging phenotypes in UK Biobank. |
| **30279459** | van der Meer D | 2018 | Brain scans from 21,297 individuals reveal the genetic architecture of hippocampal subfield volumes. |
| **30258056** | Vojinovic D | 2018 | Genome-wide association study of 23,500 individuals identifies 7 loci associated with brain ventricular volume. |
| **28924203** | Ren HY | 2017 | The common variants implicated in microstructural abnormality of first episode and drug-naive patients with schizophrenia. |
| **25607358** | Hibar DP | 2015 | Common genetic variants influence human subcortical brain structures. |
| **22504421** | Bis JC | 2012 | Common variants at 12q14 and 12q24 are associated with hippocampal volume |
| **22504418** | Ikram MA | 2012 | Common variants at 6q22 and 17q21 are associated with intracranial volume. |

#### Multi-trati colocalization (HyPrColoc) sensitivity analysis

To assess sensitivity of our result using the default settings, we repeated our analyses across a range of parameter specifications (i.e., we performed a 3-dimensional grid search with $p_c$=[0.02, 0.01, 0.005], $P_R$=[0.6, 0.7, 0.8], $P_A$=[0.6, 0.7, 0.8]). Of the 56 traits assessed (cortical eQTL and all significant associations in the KCNK2 region) and across the range of parameter settings, the traits regularly formed a single cluster of jointly colocalized traits driven by the rs1452628 variant. Occasionally, for the smallest choice of prior and largest values of threshold parameters, the "bilateral Primary intermediate ramus of the intraparietal sulcus (F.I.P.r.int.1) width" measure was removed from the cluster.

#### Summary of neuropsychiatric and cognitive phenotypes tested for genetic correlation

| Neuro trait | First author | Year | PMID | Sample size |
|---|---|---|---|---|
| **Alzheimer's disease** | Kunkle BW | 2019 | 30820047 | 63,926 |
| **Epilepsy** | Abou-Khalil B | 2018 | 30531953 | 44,889 |
| **ADHD** | Demontis D | 2019 | 30478444 | 55,374 |
| **Cognitive performance** | Lee JJ | 2018 | 30038396 | 257,828 |
| **Bipolar disorder** | Ruderfer DM | 2018 | 29906448 | 74,194 |
| **Major depressive disorder** | Wray NR | 2018 | 29700475 | 173,005 |
| **Autism spectrum disorder** | Anney RJL | 2017 | 28540026 | 15,954 |
| **Anorexia** | Duncan LE | 2017 | 28494655 | 14,477 |
| **Chronotype** | Jones SE | 2016 | 27494321 | 127,898 |
| **Generalized Anxiety disorder** | Otowa T | 2016 | 26754954 | 17,310 |
| **Parkinson's disease** | Nalls MA | 2014 | 25064009 | 108,990 |
| **Schizophrenia** | Ripke S | 2014 | 25056061 | 77,096 |

#### Biogen Biobank Team contributors

**Steering team:** Ellen Tsai, Christopher D. Whelan, Paola Bronson, David Sexton, Sally John, Heiko Runz.
**Data management team:** Eric Marshall, Mehool Patel, Saranya Duraisamy, Timothy Swan.
**Extended scientific team:** Dennis Baird, Chia-Yen Chen, Susan Eaton, Jake Gagnon, Feng Gao, Cynthia Gubbels, Yunfeng Huang, Varant Kupelian, Kejie Li, Dawei Liu, Stephanie Loomis, Helen McLaughlin, Adele Mitchell, Benjamin Sun.
